# Comparison of SARS-CoV2 N gene real-time RT-PCR targets and commercially available mastermixes

**DOI:** 10.1101/2020.04.17.047118

**Authors:** Julianne R Brown, Denise O’Sullivan, Rui PA Pereira, Alexandra S Whale, Eloise Busby, Jim Huggett, Kathryn Harris

**Affiliations:** Microbiology, Virology and Infection Prevention and Control, Great Ormond Street Hospital for Children NHS Foundation Trust, London, United Kingdom; National Measurement Laboratory at LGC, Teddington, United Kingdom; School of Biosciences & Medicine, Faculty of Health & Medical Sciences, University of Surrey

## Abstract

We aim to test four one-step RT real-time mastermix options for use in SARS-CoV2 real-time PCR, with three primer/probe assays targeting the N gene. The lower limit of detection is determined using a SARS CoV2 N gene RNA transcript dilution series (to 1 copy/µl) and verified using 74 nose and throat swabs.

The N2 assay demonstrates the most sensitive detection of SARS-Cov-2 RNA. Three of the four mastermixes performed well, with the Takara One Step PrimeScript™ III RT-PCR Kit mastermix demonstrating improved performance at the lower limit of detection.

## INTRODUCTION

The 2019/2020 pandemic of SARS-CoV2 infection, with more than 2 million confirmed cases reported globally to date (Dong *et al.* 2020), has resulted in the need for the rapid implementation of diagnostic testing centred around detection of SARS-CoV2 RNA using real-time RT PCR. The urgent need for a rapid increase in testing capacity has resulted in shortages of critical reagents, including PCR mastermixes.

This study aims to test multiple commercially available one-step RT real-time mastermix options for use in SARS-CoV2 real-time PCR testing.

## METHODS

### Test material

#### N gene transcript

An RNA transcript of the SARS-CoV2 nucleocapsid (N) gene was constructed (see supplementary methods) and quantified using the Qubit Fluorometer [Thermo Fisher].

A 10-fold dilution series of SARS-CoV2 N gene transcript was prepared from 1– 100,000 copies/µl.

#### Nose and throat swabs

Total nucleic acid was purified from 74 nose and/or throat swabs from suspected cases of SARS-CoV2 infection. Briefly, dry flocked swabs were re-suspended with 600 or 1,200 µl nuclease-free water (for single or double swabs, respectively). Total nucleic acid was purified from 200 µl swab suspension fluid using the Hamilton Nimbus (with Kingfisher Presto) and Omega Biotek Mag-Bind Viral DNA/RNA kit with 100 µl elution volume. Each specimen was spiked with Phocine Distemper Virus (PDV) cell culture isolate (cultured in vero cells) during the nucleic acid extraction as an internal positive control.

### *In silico* evaluation of primer/probe design

For the *in silico* evaluation of primer and probe coverage, 3,820 SARS-CoV-2 genome sequences were retrieved from database Global Initiative on Sharing All Influenza Data (GISAID) (Shu and McCauley, 2017) (https://www.gisaid.org/) (accessed 03.04.2020). In addition, 60 taxonomically closely related human SARS-CoV genome sequences were obtained from NCBI data, using SARS-CoV reference sequence (GenBank accession number NC_004718.3). Reduced length (<27,000 bp) and low quality sequences (≥10% unknown bases (Ns) were removed. Upon quality filtering, 3,266 SARS-CoV-2 and 61 SARS-CoV sequences were aligned using ClustalW programme in MEGA X (Kumar et al., 2018). Primer and probe coverages were calculated from the aligned sequence data, considering no mismatches.

### PCR testing

The N gene transcript dilution series, from 1– 100,000 copies/µl, was tested in duplicate using the four commercially available mastermixes in Table 1 and the three primer and probe sets in Table 2.

**Table 1.** Details of commercial mastermixes used

| Mastermix name | Manufacturer | Manufacturer product code |
| --- | --- | --- |
| One Step PrimeScript™ III RT-PCR Kit | Takara | RR600B |
| Quantifast Multiplex RT-PCR +R mastermix | Qiagen | 204956 |
| TaqPath 1-Step RT-qPCR MasterMix | ThermoFisher | A15299 |
| Taqman Fast Virus 1-step mastermix | ThermoFisher | 4444432 |

**Table 2.**
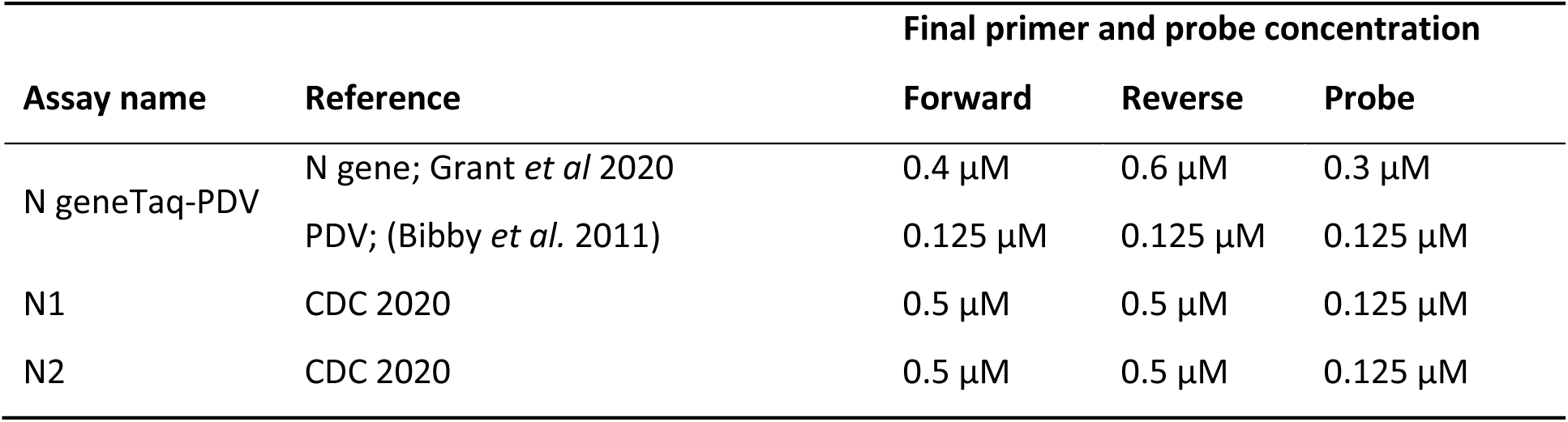
Details of primer and probes used for detection of SARS-CoV2 RNA

| Assay name | Reference | Final primer and probe concentration |  |  |
| --- | --- | --- | --- | --- |
|  |  | Forward | Reverse | Probe |
| N geneTaq-PDV | N gene; Grant *et al* 2020 | 0.4 µM | 0.6 µM | 0.3 µM |
|  | PDV; (Bibby *et al.* 2011) | 0.125 µM | 0.125 µM | 0.125 µM |
| N1 | CDC 2020 | 0.5 µM | 0.5 µM | 0.125 µM |
| N2 | CDC 2020 | 0.5 µM | 0.5 µM | 0.125 µM |

The nucleic acid purified from swabs were tested using the Takara and Qiagen mastermixes with the N geneTaq-PDV PCR assay as per Table 1 and Table 2. A negative extraction control was extracted and PCR tested alongside the swabs,

All PCR reactions were performed in 25 µl reaction volumes with 7.5 µl of RNA per reaction and run on a Quant5 thermocycler [Thermo Fisher] with manufacturer recommended fast cycling conditions and 45 cycles (Table 3).

**Table 3.**
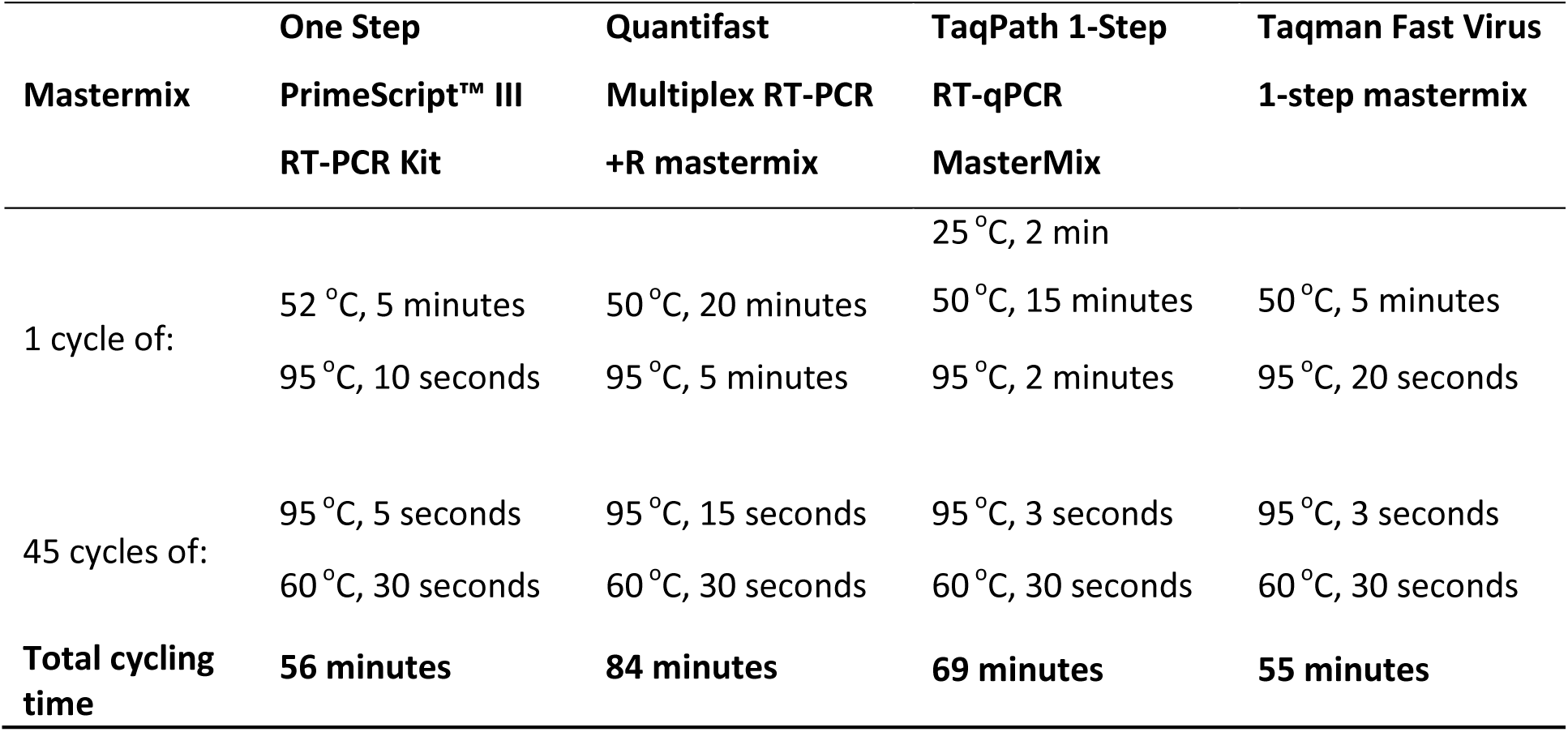
PCR cycling conditions used for each of the mastermixes tested in this study

## RESULTS

### N gene transcript dilution series

The cycle threshold (Ct) values obtained for each of the N gene transcript dilution series with each mastermix and primer and probe assay is shown in Table 4.

**Table 4.** PCR results of N gene transcript dilution series, tested in duplicate. Results are expressed as Ct values. The lower limit of detection (LLOD) is estimated to be the lowest dilution at which both (2/2) duplicates are detected. Where only one of the dilution duplicates (1/2) is detected, the LLOD is estimated to be between that and the consistently (2/2) detected dilution. Where the lowest dilution is consistently (2/2) detected with a Ct value <40, the LLOD is estimated to be less than the lowest dilution tested.

| N gene transcript (copies/μl) | | Takara | | Qiagen Quantifast | | Fast Virus | | Taqpath | |
| --- | --- | --- | --- | --- | --- | --- | --- | --- | --- |
| **N geneTaq-PDV assay** | 100,000 | 22.4 | 22.4 | 22.6 | 22.6 | 23.4 | 23.4 | 21.4 | 21.4 |
|  | 10,000 | 26.5 | 26.2 | 26.3 | 26.2 | 27.1 | 27.3 | 25.2 | 24.9 |
|  | 1000 | 29.1 | 29 | 30.2 | 30.2 | 29.8 | 29.5 | 28.5 | 28.4 |
|  | 100 | 32.6 | 32.9 | 33.4 | 33.7 | 33.3 | 33.4 | 31.7 | 31.7 |
|  | 10 | 36.2 | 37.5 | 39 | 36.5 | 39.3 | 38.8 | 35.7 | 34.8 |
|  | 1 | 39.5 | 40.2 | ND | ND | ND | ND | 38.1 | ND |
|  | *LLOD* | *1 copy/μl* |  | *10 copies/μl* |  | *10 copies/μl* |  | *1–10 copies/μl* |  |
| **CDC N1 assay** | 100,000 | 21.7 | 21.7 | 21 | 21 | 22.8 | 22.8 | 20.7 | 20.8 |
|  | 10,000 | 25.5 | 25.4 | 24.6 | 24.5 | 26.6 | 26.5 | 24.2 | 24.2 |
|  | 1000 | 28.3 | 28.4 | 28.5 | 27.9 | 29.3 | 29 | 27 | 27.1 |
|  | 100 | 31 | 31.7 | 31.1 | 31.8 | 33.1 | 32.9 | 29.9 | 31 |
|  | 10 | 35.4 | 35.6 | 36.8 | 34.8 | ND | 37 | 34.4 | 33.4 |
|  | 1 | 39.3 | 41.4 | 40.1 | ND | ND | ND | ND | 37.4 |
|  | *LLOD* | *1 copy/μl* |  | *1–10 copies/μl* |  | *10–100 copies/μl* |  | *1–10 copies/μl* |  |
| **CDC N2 assay** | 100,000 | 21.1 | 21.2 | 20.2 | 20.2 | 21.8 | 21.8 | 20.2 | 20.2 |
|  | 10,000 | 25 | 25 | 23.8 | 23.7 | 25.8 | 25.4 | 23.7 | 23.8 |
|  | 1000 | 28 | 27.7 | 26.8 | 27.1 | 28.8 | 28.6 | 26 | 27 |
|  | 100 | 31.5 | 31.5 | 30.9 | 30.7 | 32.5 | 32.5 | 30.7 | 30.3 |
|  | 10 | 34.8 | 35 | 34.7 | 35.7 | 35.4 | 35.3 | 33.7 | 34.6 |
|  | 1 | 38.1 | 38.7 | 37.6 | 36.2 | 38.9 | ND | 37.3 | 38.1 |
|  | *LLOD* | *<1 copy/μl* |  | *<1 copy/μl* |  | *1–10 copies/μl* |  | *<1 copy/μl* |  |

ND, not detected; LLOD, lower limit of detection

### Nose and throat swabs

Of the 74 swabs tested using the Takara and Qiagen mastermixes, 67/74 had concordant SARS-CoV2 PCR results (Table 5). Of the discordant results, 7/74 swabs were PCR positive with the Takara mastermix but negative with the Qiagen mastermix, with a median Ct value of 40.9 (range 38.5– 41.9). There were no specimens (0/74) positive with the Qiagen PCR but negative with Takara mastermix. All SARS-CoV2 PCR negative specimens (38/74) had detectable PDV within the expected Ct value range (data not shown). Moreover, the background fluorescence of the amplification plot was considerably lower with Takara mastermix compared to Qiagen Mastermix (Figure 1)

**Table 5.** Concordance in SARS-CoV2 PCR results for 74 swabs tested using Qiagen Quantifast Multiplex RT-PCR +R mastermix and Takara One Step PrimeScript™ III RT-PCR Kit mastermix, and in-house N geneTaq-PDV real-time RT-PCR assay. Results expressed as number of specimens.

|  | Positive by Takara | Negative by Takara |
| --- | --- | --- |
| Positive by Qiagen | 29 | 0 |
| Negative by Qiagen | 7* | 38 |

\* Median Ct 40.9 (range 38.5 – 41.9)

**Figure 1.**
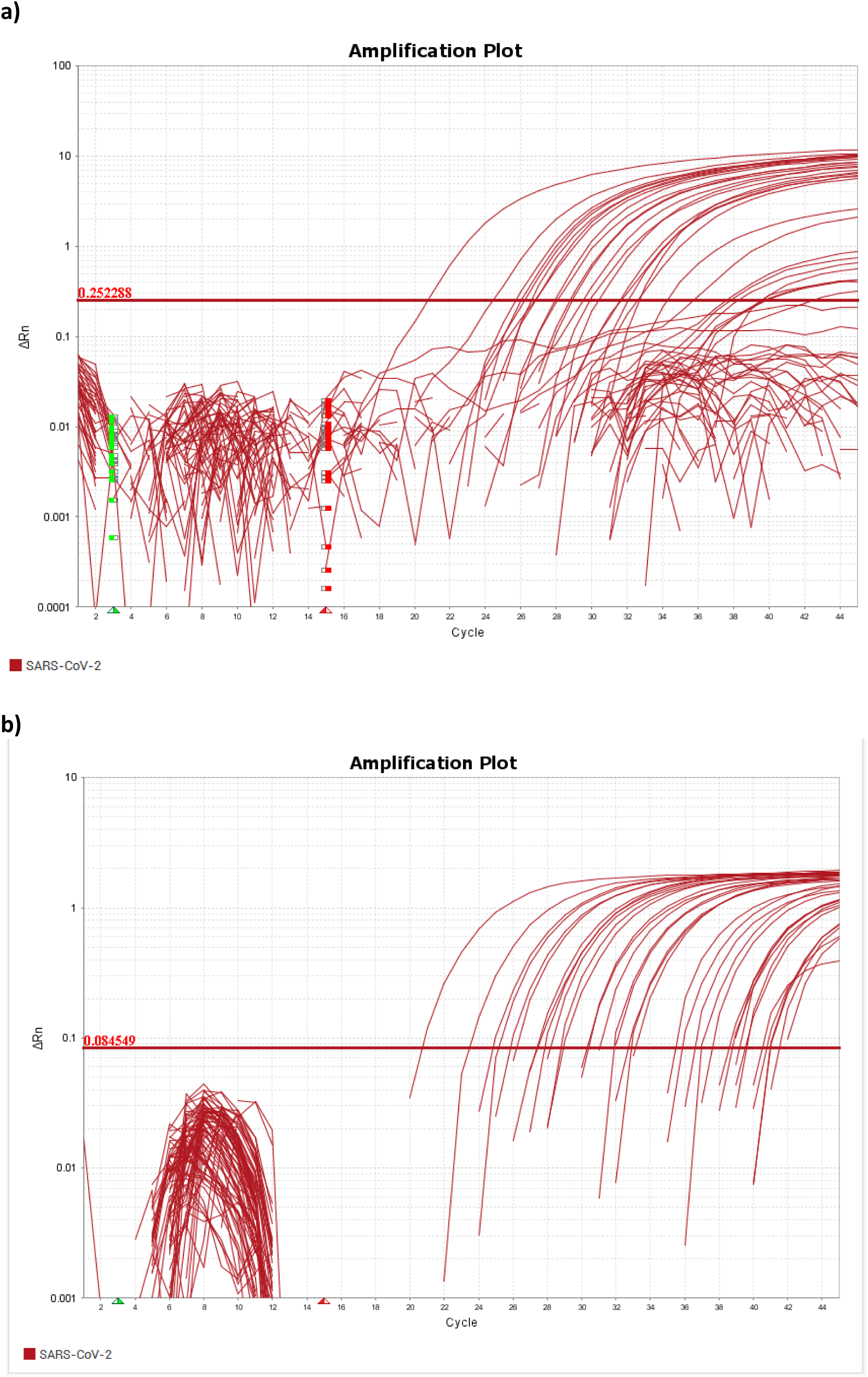
Amplification plot from 74 swabs tested using **a)** Qiagen Quantifast Multiplex RT-PCR +R mastermix and **b)** Takara One Step PrimeScript™ III RT-PCR Kit mastermix, and in-house N geneTaq-PDV real-time RT-PCR assay on Quant5 thermocycler.

### *In silico* evaluation of primer/probe design

All three primer/probe sets tested in this study (N geneTaq, N1 and N2) are identical to >99% of SARS-CoV2 sequences available up to 3^rd^ April 2020 (see Supplementary Results). N geneTaq primers (Grant *et al.* 2020) were also identical to SARS-CoV sequences, with mismatches in 4/22 positions in the probe. The N1 primers and probe had mismatches to SARS-CoV in 2–7 positions in each primer and probe. The N2 forward primer is identical to SARS-CoV, with mismatches in 2–5 positions in each the reverse primer and probe.

## CONCLUSIONS

We demonstrate that, of the RT-PCR assays targeting SARS-CoV2 N gene RNA tested in this study, the N2 primers and probe (CDC 2020) show the lowest limit of detection across all four of the mastermixes tested (<1 copy/µl with Takara, Qiagen and TaqPath; 1–10 copies/µl with Fast Virus).

Our data shows that the Takara One Step PrimeScript™ III RT-PCR Kit mastermix has improved performance at the lower limit of detection compared to the other mastermixes tested in this study, evidenced by a lower limit of detection of the N gene RNA transcript across all N gene targets (LLOD <1–10 copies/µl) and the additional detection of 7/74 swabs with our in-house N geneTaq-PDV assay. Taqman Fast Virus 1-step mastermix is the most poorly performing of the mastermixes tested, with a LLOD of 1–100 copies/µl across N gene targets tested.

All three primer/probe sets (N geneTaq, N1 and N2) have good homology to sequenced strains of SARS-CoV2 so are likely to detect circulating virus. Based on homology to SARS-CoV sequences, N geneTaq primers and probe (Grant *et al.* 2020) are the most likely to show some cross-reactivity with SARS-CoV virus but requires *in vitro* testing to verify, as the mismatches in the probe may be sufficient to prevent cross-amplification. Possible cross-reactivity with SARS-CoV is not of clinical concern at this time. No cross-reactivity is anticipated with other circulating human betacoronaviruses.

Based on the data generated in this study we recommend that for optimum detection of SARS CoV-2 RNA from nose and throat swabs the N2 primers and probe assay is used with any of the following mastermixes; Takara, Qiagen and TaqPath. Alternatively we recommend the N geneTaq (Grant *et al* 2020) or the N1 assays are used with Takara mastermix.

The data presented here provides evidence for the recommendation of mastermixes for detection of SARS-CoV2 RNA targeting the N gene and suggests adequate alternatives in the event of supply chain issues.

## Supporting information

Supplementary Results

## REFERENCES

Bibby DF, McElarney I, Breuer J and Clark DA (2011). Comparative Evaluation of the Seegene Seeplex RV15 and Real-Time PCR for Respiratory Virus Detection, Journal of Medical Virology, 83: 1469–1475.

CDC (2020). Accessed online 14/04/20 https://www.cdc.gov/coronavirus/2019-ncov/lab/rt-pcr-panel-primer-probes.html

Dong E, Du H and Gardner L (2020). An interactive web-based dashboard to track COVID-19 in real time, The Lnacet Infectious Diseases, DOI: https://doi.org/10.1016/S1473-3099(20)30120-1

Grant PR, Turner MA, Shin GY, Nastouli E and Levett LJ (2020). Extraction-free COVID-19 (SARS-CoV-2) diagnosis by RT-PCR to increase capacity for national testing programmes during a pandemic, bioRxiv, doi: https://doi.org/10.1101/2020.04.06.028316

Kumar, S., Stecher, G., Li, M., Knyaz, C., and Tamura, K. (2018). MEGA X: Molecular Evolutionary Genetics Analysis across Computing Platforms. Mol Biol Evol 35, 1547–1549. doi:10.1093/molbev/msy096.

Shu, Y., and McCauley, J. (2017). GISAID: Global initiative on sharing all influenza data –from vision to reality. Euro Surveill 22. doi:10.2807/1560-7917.ES.2017.22.13.30494.

