## Supplementary Methods for "Comparison of SARS-CoV2 N gene real-time RT-PCR targets and commercially available mastermixes"

### Preparation of RNA Transcript:

A genetic construct containing the partial sequence of the SARS-CoV2 nucleocapsid (N) gene (location: SARS-CoV-2 genome region 28274-29239 MN908974.3) was synthesised and inserted into a pUC19 plasmid vector (New England Biolabs) by Eurofins MWG Operon. The plasmid (41.2 ng) was linearised using the restriction enzyme *BamHI* (New England Biolabs) (0.4 units/**µL**), CutSmart Buffer and nuclease free water (Ambion) in a reaction volume of 50 µL were incubated at 37$^{o}$C for 1 hour. The linearised plasmid was purified using the QIAquick PCR purification kit (Qiagen) according to manufacturer guidelines. The plasmid was analysed using the Agilent 2100 BioAnalyzer with DNA 7500 series II kit (Agilent) to confirm the expected fragment length of 3450 bp and quantified using the High Sensitivity dsDNA assay with the Qubit® 2.0 Fluorometer (Thermo Fisher).

An RNA transcript was prepared by *in vitro* transcription using the T7 RNA polymerase (MEGAScript T7 kit, Ambion Life Technolgies) according to manufacturer's guidelines and subsequent purification using the RNeasy Mini Kit for RNA clean up (Qiagen) which included an on column DNase I step (RNase-free DNase set, Qiagen). Dilutions of the *in vitro* transcribed RNA were prepared in RNA Storage solution (Thermo Fisher) and aliquots stored at -80$^{o}$C for further use. This prepared material was analysed using the Agilent 2100 BioAnalyzer with RNA 6000 kit (Agilent) and quantified using the High Sensitivity RNA assay with the Qubit® 2.0 Fluorometer (Thermo Fisher).  Dilutions of this were prepared in nuclease-free water.

Sequence of the RNA Transcript

| Partial N gene (+3G from T7 promoter and BamHI sequence from anti-sense DNA strand) |
|---|
| GGGAUGUCUGAUAAUGGACCCCAAAAUCAGCGAAAUGCACCCCGCAUUACGUUUGGUGGACCCUCAGAU<br>UCAACUGGCAGUAACCAGAAUGGAGAACGCAGUGGGGCGCGAUCAAAACAACGUCGGCCCCAAGGUUUA<br>CCCAAUAAUACUGCGUCUUGGUUCACCGCUCUCACUCAACAUGGCAAGGAAGACCUUAAAUUCCCUCGA<br>GGACAAGGCGUUCCAAUUAACACCAAUAGCAGUCCAGAUGACCAAAUUGGCUACUACCGAAGAGCUACC<br>AGACGAAUUCGUGGUGGUGACGGUAAAAUGAAAGAUCUCAGUCCAAGAUGGUAUUUCUACUACCUAGG<br>AACUGGGCCAGAAGCUGGACUUCCCUAUGGUGCUAACAAAGACGGCAUCAUAUGGGUUGCAACUGAGG<br>GAGCCUUGAAUACACCAAAAGAUCACAUUGGCACCCGCAAUCCUGCUAACAAUGCUGCAAUCGUGCUAC<br>AACUUCCUCAAGGAACAACAUUGCCAAAAGGCUUCUACGCAGAAGGGAGCAGAGGCGGCAGUCAAGCCU<br>CUUCUCGUUCCUCAUCACGUAGUCGCAACAGUUCAAGAAAUUCAACUCCAGGCAGCAGUAGGGGAACUU<br>CUCCUGCUAGAAUGGCUGGCAAUGGCGGUGAUGCUGCUCUUGCUUUGCUGCUGCUUGACAGAUUGAAC<br>CAGCUUGAGAGCAAAAUGUCUGGUAAAGGCCAACAACAACAAGGCCAAACUGUCACUAAGAAAUCUGCU<br>GCUGAGGCUUCUAAGAAGCCUCGGCAAAAACGUACUGCCACUAAAGCAUACAAUGUAACACAAGCUUUC<br>GGCAGACGUGGUCCAGAACAAACCCAAGGAAAUUUUGGGGACCAGGAACUAAUCAGACAAGGAACUGAU<br>UACAAACAUUGGCCGCAAAUUGCACAAUUUGCCCCCAGCGCUUCAGCGUUCUUCGGAAUGUCGCGCAUU<br>GGCAUGCCUAG |
